# An accelerated thrombosis model for computational fluid dynamics simulations in rotary blood pumps

**DOI:** 10.1101/2021.08.30.458209

**Authors:** Christopher Blum, Sascha Groß-Hardt, Ulrich Steinseifer, Michael Neidlin

**Author notes:** **Correspondence:** Name: Michael Neidlin, Address: Institute of Applied Medical Engineering, Pauwelsstraße 20, 52074 Aachen, Germany.

## Abstract

**Purpose:** Thrombosis is one of the major complications in blood-carrying medical devices and a better understanding to influence design of such devices is desirable. Over the past years many computational models of thrombosis have been developed. However, open questions remain about the applicability and implementation within a pump development process. The aim of the study was to develop and test a computationally efficient model for thrombus risk prediction in rotary blood pumps.

**Methods:** We used a two-stage approach to calculate thrombus risk. At the first stage, the velocity and pressure fields were computed by computational fluid dynamic (CFD) simulations. At the second stage, platelet activation by mechanical and chemical stimuli was determined through species transport with an Eulerian approach. The model was implemented in ANSYS CFX and compared with existing clinical data on thrombus deposition within the HeartMate II.

**Results:** Our model shows good correlation (R^2^>0.94) with clinical data and identifies the bearing and outlet stator region of the HeartMate II as the location most prone to thrombus formation. The calculation of platelet activation requires an additional 10-20 core hours of computation time.

**Discussion:** The concentration of activated platelets can be used as a surrogate marker to determine risk regions of thrombus deposition in a blood pump. Model expansion, e.g. by including more chemical species can easily be performed. We make our model openly available by implementing it for the FDA benchmark blood pump.

**Declarations:** *Funding:* This research received no specific grant from any funding agency in the public, commercial, or not-for-profit sectors. Open access funding enabled and organized by Projekt DEAL.

*Conflict of interest:* All of the authors have nothing to disclose.

*Availability of data and material:* The raw data can be retrieved by request from the authors.

*Code availability:* The implementation of the thrombus model in the FDA benchmark blood pump geometry is available on https://doi.org/10.5281/zenodo.5116063.

*Authors’ contributions:* All authors contributed to the study conception and design. CB developed the numerical model, performed the simulations, gathered, analysed and discussed the results. SGH, MN and US were involved in the analysis and discussion of the results. MN supervised the project. MN and CB wrote the manuscript based on the input of all co-authors. All co-authors read and approved the final version of the manuscript.

## Introduction

Rotary blood pumps such as ventricular assist devices (VADs) and extracorporeal pumps are used in long-term and short-term mechanical circulatory support therapies for patients with acute and chronic heart failure [1, 2]. Despite tremendous improvements in patient survival and overall quality of life over the last decades, one of the major device related complications is the formation of thrombi (blood clots) and the associated risk for cardioembolic stroke [3]. The causes for thrombus formation are multifactorial involving the molecular interplay of proteins and enzymes in the coagulation cascade, non-physiological blood flow conditions and interactions with foreign surfaces [4]. Ultimately the aforementioned factors enhance the hemostatic response by chronically activating platelets, the initiators of blood clots [5].

Computational modeling can help to understand the underlying complexity of thrombus formation involving a wide range of mathematical models of thrombosis spanned across different spatial and temporal scales [6]. Considering model application in blood-carrying medical devices in general and rotary blood pumps in particular poses additional requirements towards thrombosis models. They should be implementable within computational fluid dynamics (CFD) solvers, the gold-standard to determine blood flow characteristics, such as velocity, pressure and shear stress distributions, and computable for arbitrary geometries. In addition, the computational costs should be kept at an appropriate level, as already the flow physics within rotary blood pumps are highly intricate including turbulent flow structures, complex geometries and high rotational speeds. Optimally, such a model should be compatible with commercial CFD software to allow a wider adoption throughout the scientific community. Several approaches that serve as good candidates exist in the literature. Many studies in the past adopted the idea of a platelet activation state (PAS) [7, 8] that determines stress accumulation of platelet flow paths either through Eulerian or Lagrangian approaches. In consequence, each device exhibits a “thrombogenicity fingerprint” that can then be compared between different devices. This approach is similar to the hemolysis models existing in the literature [9]. Despite being computationally cheap, it might be difficult to obtain spatially resolved thrombus deposition within the domain, although there exist recent approaches that try to achieve this [10].

The thrombosis model presented in Taylor et al. [11] uses CFD with species transport of non-activated platelets, activated platelets and adenosine diphosphate (ADP) by means of convection diffusion equations. In addition, a Brinkman term is added to the momentum equations of the flow solution to assign areas specified as thrombus with a velocity approaching zero. This achieves a coupling of the flow solution and the thrombus growth. The model has so far been applied to a simple backward facing step geometry and has been validated with time-resolved in vitro experiments [12, 13]. For this geometry, experimental thresholds of wall shear stresses, which control the growth and detachment of thrombus areas, have been determined. However, these values could be dependent on the geometry and flow conditions. To the best of our knowledge, the model has not been applied in more complex flow geometries such as blood pumps. Furthermore, the model has been developed for laminar flows and resolves the entire time scale until a stable thrombus is reached. This proves to be impractical, especially for the simulation of rotating cardiovascular devices where small time steps are required for the necessary accuracy. Hence, it would be too computationally expensive to simulate the entire physical time of around 30 min which is necessary for the formation of a stable thrombus.

Another thrombosis model is presented in Wu et al. [14]. This model follows a similar approach as the Taylor model, but includes 10 different biological and chemical species to allow a finer description of thrombosis including activation, deposition, accumulation and clearing of platelets. For the correct use of the model, 3 parameters such as the deposition rate of platelets for the materials that come into contact with the blood must be determined experimentally. The model has already been applied in [15] to a cardiovascular device (HeartMate II VAD). However, the simulations were still quite computationally intensive (5 days using 40 cores with 2.4 GHz) and had limited predictive power when compared to clinical studies. More specifically, thrombus formation was predicted only in one region (stator) and not in other known/common regions such as the blade area of the impeller or further downstream locations such as the bearing and diffusor, see Figure 5 in [15] or Figure 5 in [16].

Besides these models there exist many studies using the flow field results to identify regions prone to thrombus formation such as recirculation regions [17, 18], low velocities and low washout [19] or high residence times [20]. However, all of these studies do not model thrombosis (or parts of thrombosis) explicitly. In addition, there are other approaches using multiscale simulations [21] and other computational methods besides CFD [22]. The interested reader is referred to the extensive overviews existing in the literature [6, 23].

In summary, there is still a lack of computational models capable of predicting locally resolved thrombus formation in complex geometries with reasonable computational effort.

The aim of this study is to develop such a model for rotary blood pumps focusing on low computational costs and ease of implementation in a state-of-the-art CFD solver. Moreover, information about parameter sensitivity of all mechanisms important for the computational modeler and a ready-to-use setup are provided.

## Methods

### Accelerated thrombosis model

The accelerated thrombosis model is closely based on the thrombosis model presented by Taylor et al. [11] with several reductions and adjustments to save computational time. A graphical overview of our model is shown in the Supplementary Figure S1. Detailed information about the Taylor model can be taken from the original paper.

In brief, our approach uses three convection-diffusion equations describing the species transport of non-activated platelets (NP), activated plates (AP) and adenosine diphosphate (ADP). Platelet activation (change from NP to AP) occurs either with mechanical cues through a shear stress based power law model [8], equation 1, or with chemical cues through a threshold based activation triggered by the ADP concentration, equation 2.

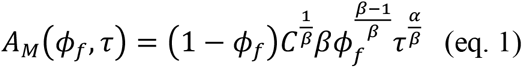

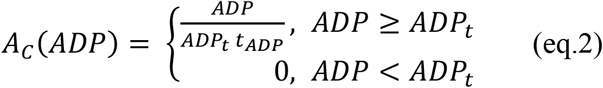

*A*_*M*_ and *A*_*C*_ are the mechanical and chemical activates rates, respectively. *φ*_*f*_ is the ratio of activated platelets to total platelet count. *α*, *β* and *C* are the power law model parameters taken from [8]. *ADP*_*t*_ is the threshold for chemical platelet activation and *t*_*ADP*_ is the characteristic time of the platelet activation rate. Finally, *τ* is the scalar shear stress calculated according equation 3 with *σ*_*ij*_ as the individual components of the shear stress tensor.

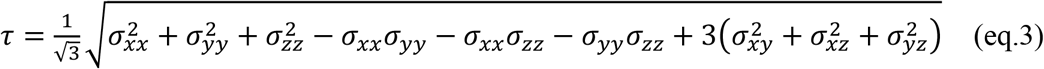

Both activation cues are summed up to determine the rate of platelet activation that is included in the according convection-diffusion equations as a source component. The AP species is considered as the model output and represents the thrombus risk.

In addition, the implementation of the laminar shear stress tensor presented so far was also compared with a turbulent implementation. To obtain a turbulent description of the shear stress tensor, the laminar part was extended with the Reynolds stress components [24]. Equations 4 and 5 show the definition of the total shear stress tensor with the laminar part on the left side and the turbulent Reynolds stress part on the right side of the minus sign.

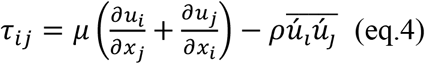

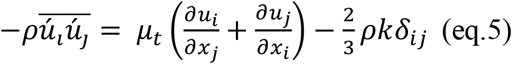

In these equations *μ* is the dynamic viscosity, *μ*_*t*_ the eddy viscosity, *ρ* the density of the fluid, *k* the turbulent kinetic energy, *δ*_*ij*_ the Kronecker delta, *u*_*i*_ the velocity components and *x*_*i*_ the spatial components.

In comparison to the Taylor model we did not include a sink in the momentum equations through a Brinkman term which reduces the velocity of regions with high AP concentrations. Moreover, the accelerated thrombosis model uses an uncoupled approach by first determining the velocity field within the pump through a (steady state or time-averaged) CFD simulation and then, in a second calculation, resolving the species transport based on the output of the first simulation. The implemented model in the FDA benchmark blood pump [25] together with a tutorial on how to run the simulations can be found at https://doi.org/10.5281/zenodo.5116063.

An overview of all parameters used in the accelerated thrombosis model are given in Table 1. In addition to the formula symbols, the definition and the according equation, this table also contains information on how the values of the parameters were determined and whether the parameters were part of the parameter study presented later on. Finally the values of the parameters are also shown. The sources of the parameters can be found in Taylor et al. [11]. With the underlying information for R_ADP_ found in [26] and for ADP_t_ found in [27]. In the column sensitivity assessment, “not complete” means that this variable was varied but no complete sensitivity analysis was carried out.

**Table 1:** Overview of parameters involved in accelerated thrombosis model

| Variable | Definition | Equation | Sensitivity<br>assessment | Determination | Initial Value |
| --- | --- | --- | --- | --- | --- |
| $D_a$ | Diffusion coefficient for activated platelets | Convection-diffusion Activated platelets | no | mechanistic | $1.58e-11$<br>[m<sup>2</sup> s<sup>-1</sup>] |
| $D_n$ | Diffusion coefficient for non-activated platelets | Convection-diffusion non-activated platelets | no | mechanistic | $1.58e-11$<br>[m<sup>2</sup> s<sup>-1</sup>] |
| $D_{ADP}$ | Diffusion coefficient for ADP | Convection-diffusion ADP | no | mechanistic | $1e-20$ [m<sup>2</sup> s<sup>-1</sup>] |
| $R_{ADP}$ | Amount of ADP released during one platelet activation | Chemical activation | no | empirical | $3e-17$ [mol] |
| $ADP_t$ | Threshold value for activation of platelets | | not complete | assumed | $2e-3$ [mol m<sup>-3</sup>] |
| $t_{ADP}$ | Characteristic time of platelet activation rate | | not complete | assumed | 1 [s] |
| C | coefficients of power law model | Mechanical activation | yes | empirical | 1.4854e-7 |
| $\alpha$ | | | yes | empirical | 1.4854 |
| $\beta$ | | | yes | empirical | 1.4401 |
| $\tau$ | Scalar shear stress | | yes | CFD | - |
| $\phi_a$ | Concentration of activated platelets | Initial condition | yes | assumed | 25e12 [m<sup>-3</sup>] |
| $\phi_b$ | Concentration of non-activated platelets | Initial condition | yes | assumed | 475e12 [m<sup>-3</sup>] |

### Geometries, mesh and simulation parameters

A reverse engineered version of the HeartMate II (HM2) geometry, an axial flow ventricular assist device (VAD), was used. The pump geometry was meshed with ANSYS 2021 R1 Meshing tool (ANSYS Inc., Canonsburg, USA using unstructured tetrahedral elements and prism layers to assure a y+ value <1. The bearings were modelled as hemispherical domains of 0.05 mm thickness and a bearing shell diameter of 3 mm, see Supplementary Figure S2, and meshed using the sweep mesh method with a segmentation of 20. Mesh independence studies resulted in mesh element numbers of approx. 12 million elements for the HM2 geometry. The non-Newtonian blood model by Ballyk et al. [28] with a density of ρ = 1056.4 kg m^-3 (hematocrit level of 44%) was used. The k-ω Shear Stress Transport model was taken as the turbulence model. For the steady-state initialization simulations, the auto-time step method was used, and the stabilization of the pressure difference between the inlet and outlet of the simulation was used as the termination criterion. For the simulations of the accelerated thrombosis model, a physical time step of one second was used and the stabilization of the volume-averaged AP concentration in each simulation domain was used as the termination criterion.

The computational mesh of the pump is shown in Figure 1 with blue as the rotating parts and grey as the non-rotating parts. The baseline simulations for the HM2 were performed at 9000 rpm and a flow of 2 l/min. ANSYS CFX was used as the CFD solver.

**Figure 1:**
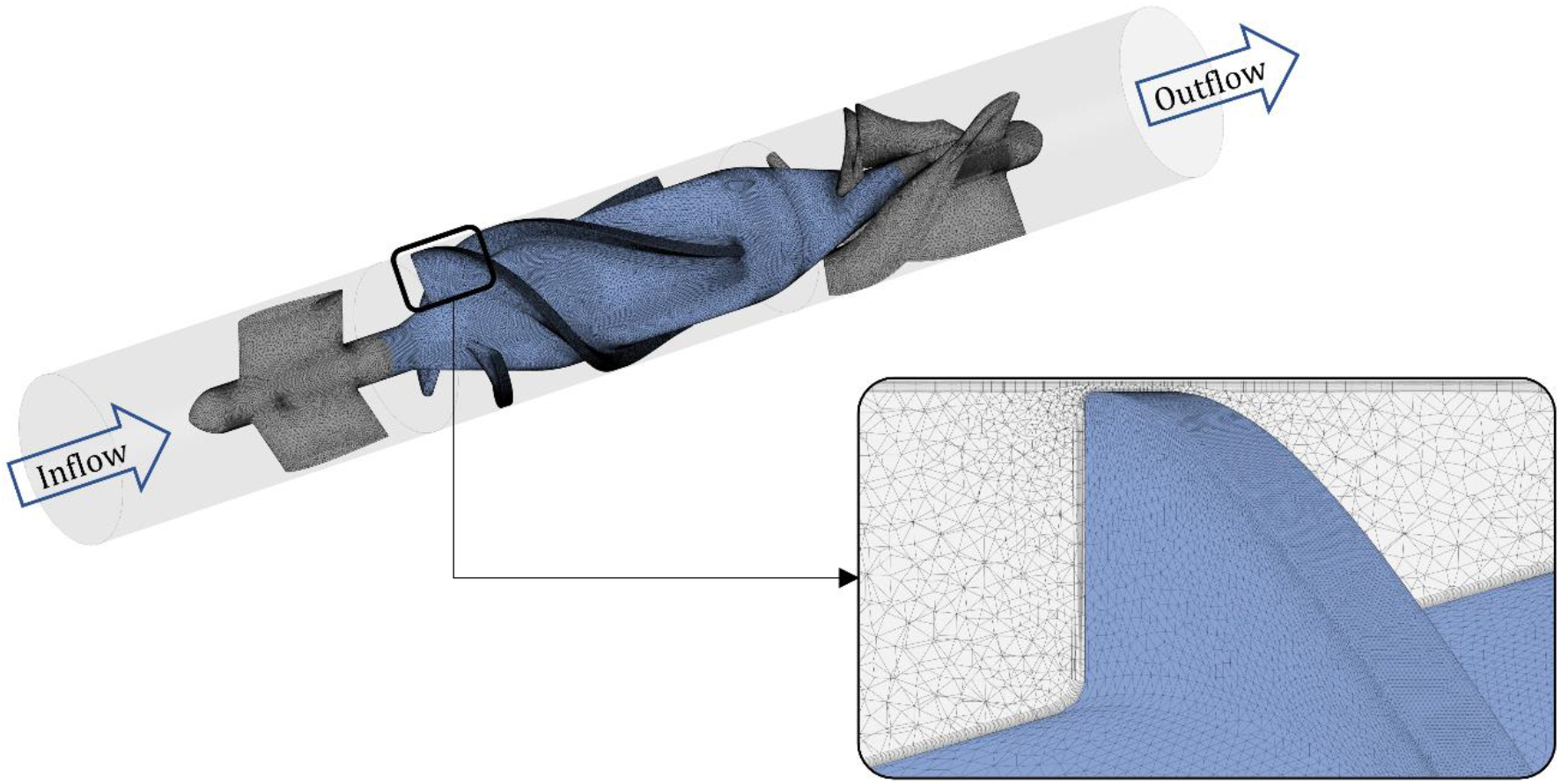
HM2 pump with flow direction and meshing details.

### Comparison with clinical data

In order to compare the computational model with in vivo data on thrombus occurrence in a blood pump, the study by Rowlands et al. [16] was used. In this study, a retrospective analysis of 29 explanted HM2 was performed to identify the frequency, composition and localization of macroscopic thrombi. The locations used in [16] were named as following: Region 1 – Inflow straightener, region 2 – Fore bearing, region 3 – Impeller Blade-Blade, region 4 - Outlet bearing/Stator. It has to be noted, that in [16] another region upstream of the inflow straightener, the fore connector seam, was considered. As this region was not present in our HM2 geometry, it could not be included in the analysis. As there was no information on the respective operating points from the explanted pumps, the baseline 9000 rpm at 2 l/min was assumed and later expanded for more speeds and pump flows to cover a broader operating range. This also corresponds to the operating point chosen in the study [15].

### Operating point and sensitivity analysis

An operating point analysis for the HM2 was performed to determine the influence of speed variation (7000-11000 rpm) and the flow rate (1 l/min-4 l/min) on the model output. Sensitivity analysis was performed for the shear stress power law parameters C, α and β (+/− 10% variation and +/−20% for the most sensitive parameter) and the initial values of background activation φ_a_ and φ_b_ (ratio between activated and non-activated platelets of 1%, 5%, 10% and 20%). As preliminary simulations did not deliver a high contribution of chemical activation, the parameters ADP_t_ and t_ADP_ were only adjusted once to identify their influence on the results. Lastly, the influence of the Reynolds stresses (latter part of the right side of equation 4) on the platelet activation was evaluated.

## Results

### Model results and comparison with clinical data

The model results with two visualizations of the AP variable are shown in Figure 2. For better readability the AP species was scaled such that the per mille increase in relation to the initial value of 25e12[m^-3] was determined and this scaling was kept consistent throughout the manuscript. In the upper section, the geometry is described by blue dots and blood volumes of scaled AP concentration above 5.7 are shown in red. Small volumes of elevated scaled AP concentration exist behind the first stator and along the blade edges. Larger volumes can be seen in the rear stator. In the lower section, the scaled value of the AP variable is displayed on the surface by a contour plot. The scale is chosen such that differences on the surface are clearly visible. To ensure comparability of the results, the same scale is used throughout the manuscript unless otherwise specified. On the surface, the areas of the trailing edges of the front stator, parts of the blade passage and especially the areas after the bladed area can be identified with high AP values.

**Figure 2:**
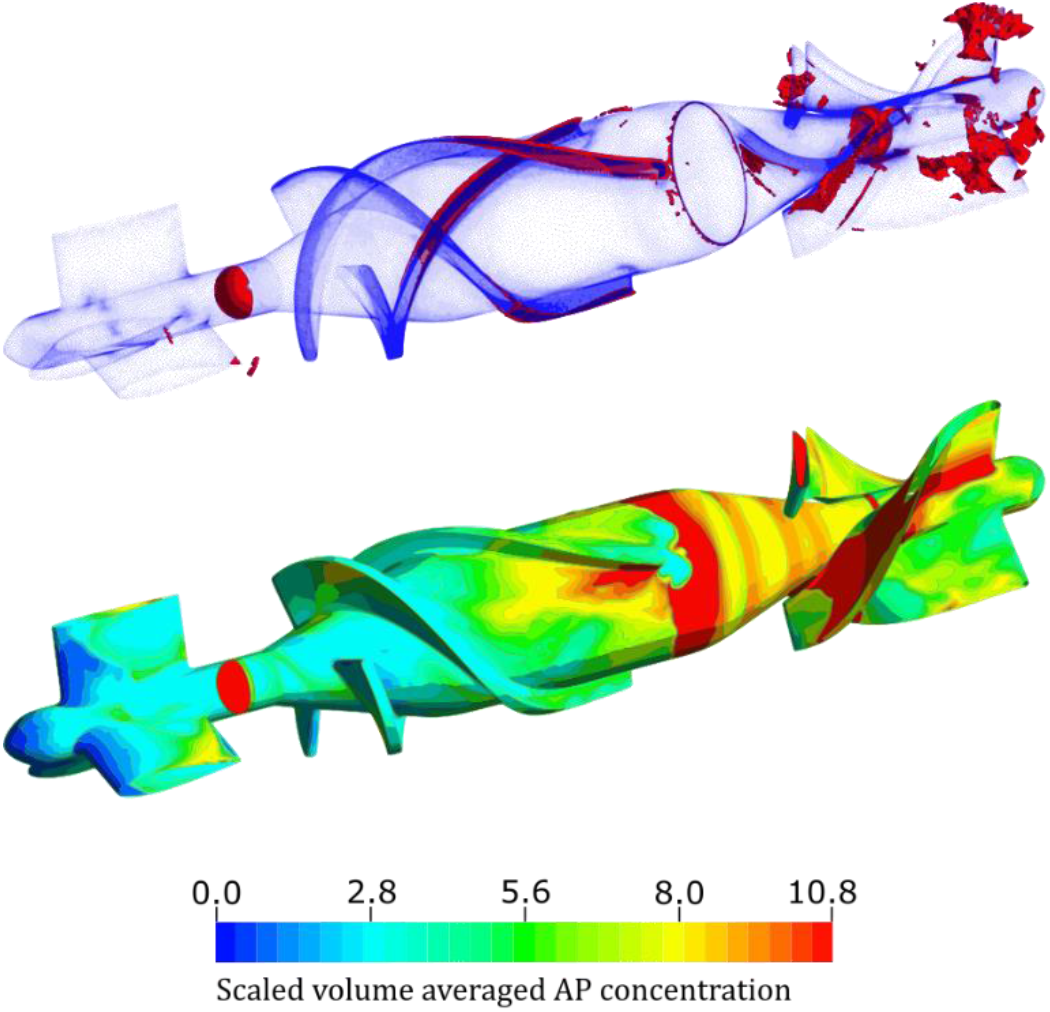
Scaled activated platelet (AP) concentration in HeartMate II geometry with 9000rpm and 2L/min

Figure 3a) shows the volume averaged scaled AP concentration in the respective fluid domains and Figure 3b) shows the frequency of the thrombi as identified from the study [16]. An excellent correlation between model output and observed thrombi exists, (R=0.99).

**Figure 3:**
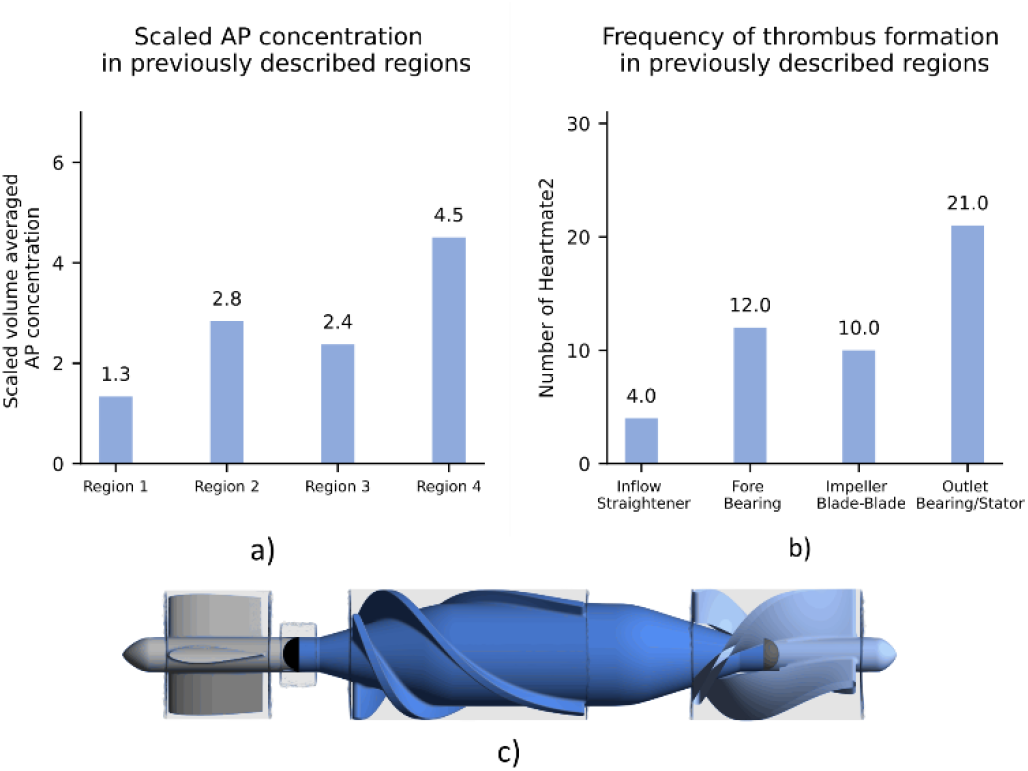
Model output and comparison with clinical data. a) Volume averaged and scaled AP in four computational domains, b) – Thrombus occurrence observed in Rowlands et al., c)-Regions in pump geometry

### Operating point analysis

The influence of speed variation and flow rate on the AP concentration is shown on the left side of Figure 4 a). The data shown in black corresponds to the previously considered operating point of 9000 revolutions per minute and 2 l/min. As the speed increases, the concentration of AP in each volume also increases. The level of AP concentration in region 3 approaches the concentration level of region 2 as the speed increases, meaning that the platelet activation potential of the bladed impeller area approaches the potential of the bearing region. In general, there is an approximately linear relationship between speed and AP concentration.

**Figure 4:**
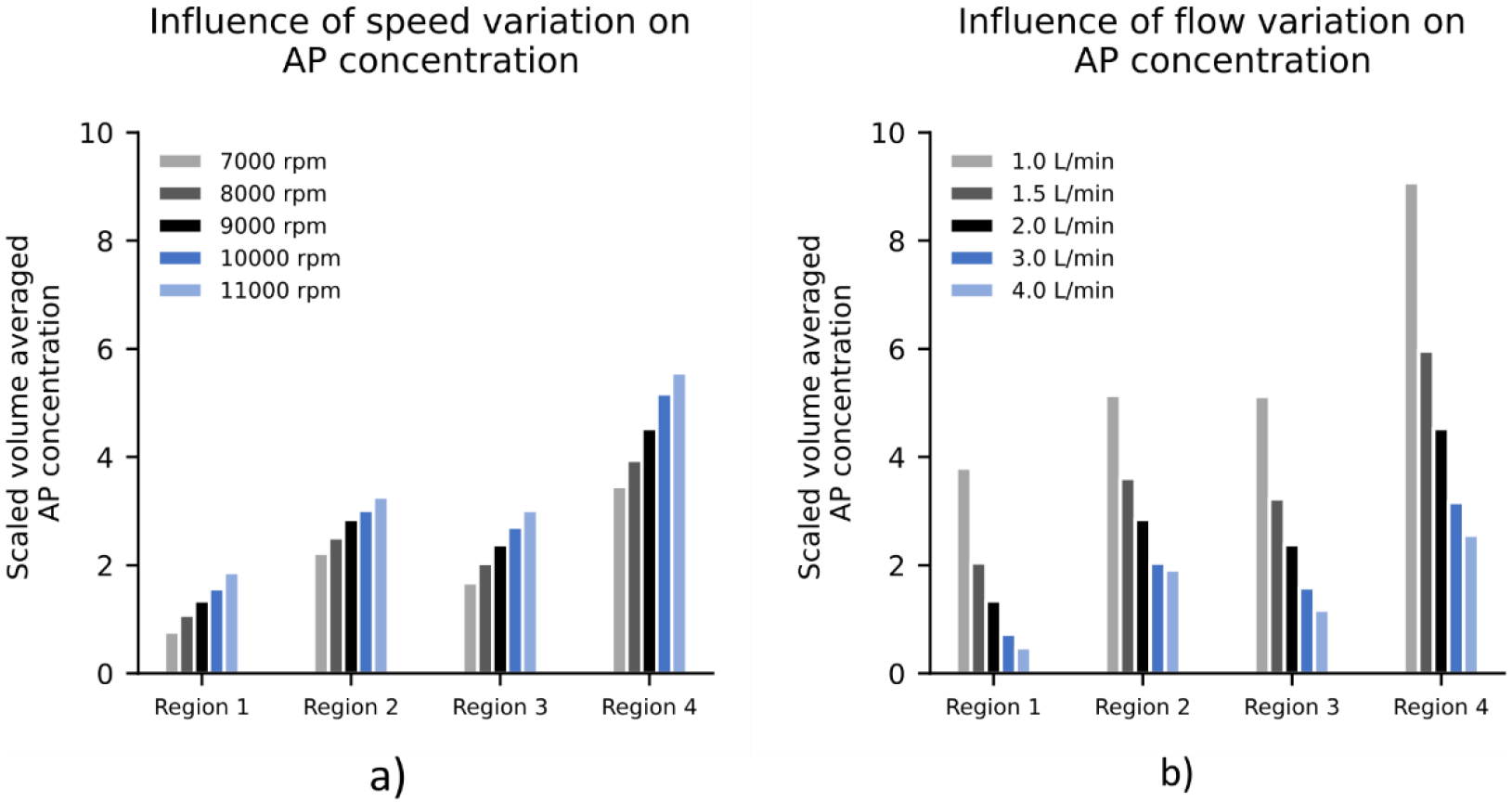
Scaled volume averaged AP concentration in HM2 for rpms between 7000-11000 rpm (a) and 1-4 l/min (b) for the regions 1-4.

Figure 4 b) shows that low flow operating points have substantially higher AP concentration values than high flow operating points indicating an inversely proportional relationship between flow rate and platelet activation.

In the most unfavorable case of 1 l/min, a substantially higher concentration of AP can already be seen in region 1 (inflow straightener) and in region 4 (outlet bearing/stator), which exhibits the highest AP concentration of all regions.

Figure 5a) shows the surface distribution of the AP variable for 1, 2 and 4 l/min flow at 9000 rpm and Figure 5b) the volume renderings of scaled AP concentrations higher than 5.7. The different flow conditions, which result from the different operating points, also result in different distributions of the AP concentration on the surface. In the case of 1 l/min almost the entire wake regions of the impeller has high AP concentration values. A comparison between the 2 l/min scenario and the 4 l/min scenario exhibits differences in AP concentration at the trailing edge of the first stator. In addition, the shape of the AP distribution in the bladed passage changes and the concentration of AP in the rear area is lower in the 4 l/min than in the 2 l/min case. A common feature of all simulations is that the two bearing areas are always indicated with a high AP concentration.

**Figure 5:**
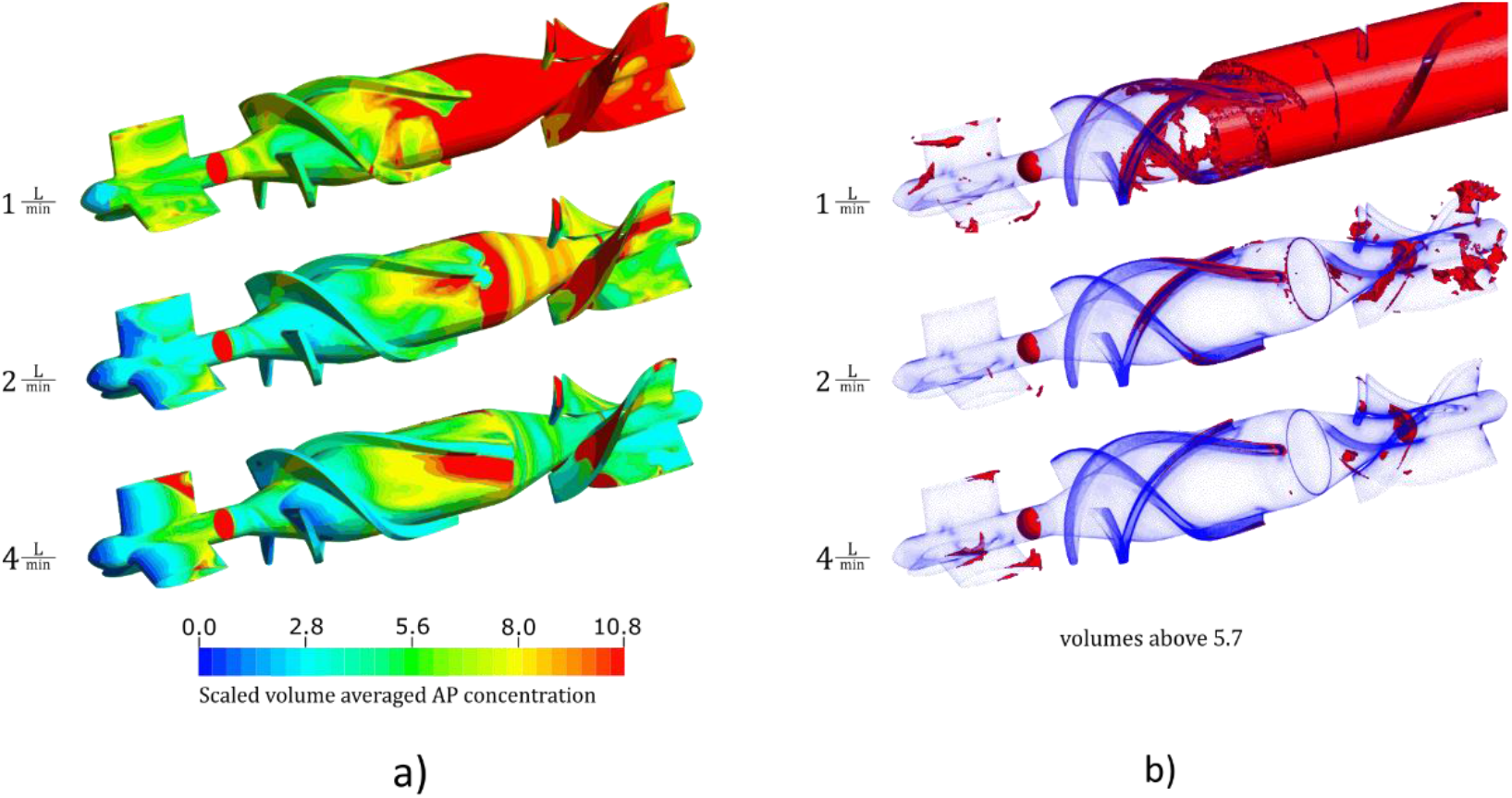
Comparison of different simulations with varying flow from 1 to 4 L/min at 9000 rpm using contour plots (a) and (b) volume renderings of the scaled variable AP

The correlation between the different operating points and the clinical data is shown in the Supplementary Figure S3. The lowest R^2^ for all scenarios was 0.94.

### Sensitivity analysis

#### Shear stress power law model

The percentage deviation of the AP concentration after parameter variation of *α*, *β* and *C* for the baseline operating condition is shown in Figure 6a). The front and back bearings are presented as separate regions. A change in the parameter *α* results in a high percentage deviation of the AP concentration, especially in the bearing areas. The parameters *β* and *C* do not seem to have a major influence on the results. In Figure 6b), the correlation between simulation and clinical data for the different perturbations of *α* is shown. Although the lines differ in intercept and slope, they all show a good correlation accuracy with the study [16]. As observable in the Supplementary Figure S4, changes of *α* influence the absolute values of AP concentration, however the spatial distribution of high and low AP concentrations remains the same.

**Figure 6:**
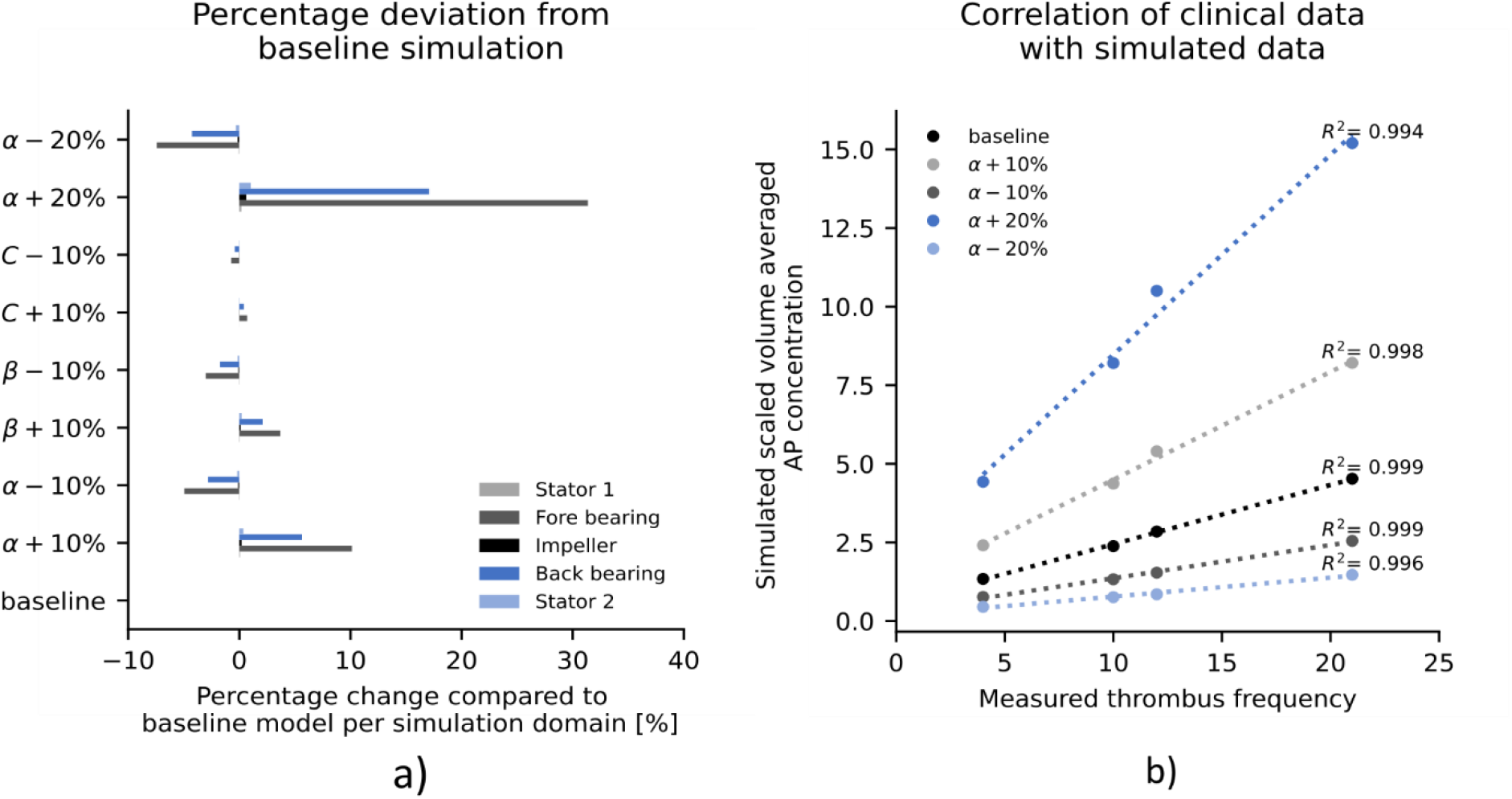
Perturbation of the parameters α, β and C of the power law model and correlation with the clinical study from Rowlands et al.

#### Influence of turbulence modeling

Whether the scalar shear stress is obtained with the turbulent stress tensor or with the laminar stress tensor also has an influence on the mechanical activation and the results are shown in Figure 7. Turbulent stresses increase platelet activation, especially in the region downstream of the impeller blades (region 4). Contour plots of the AP concentration throughout the pump are found in the Supplementary Figure S5. The qualitative behavior of the AP concentration throughout the different volumes does not change.

**Figure 7:**
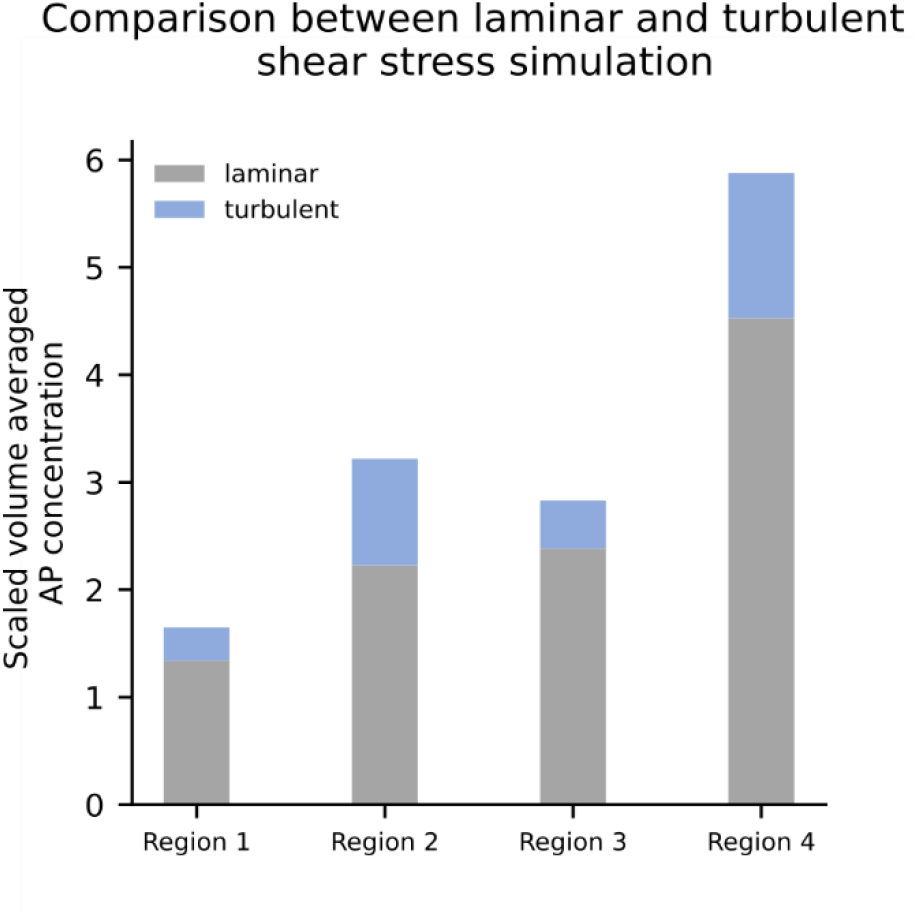
Influence of laminar or turbulent shear stress modeling on the AP distribution

#### Influence of different background activation levels

The influence of the ratio of activated platelets to non-activated platelets (ratios of 1%, 5%, 10% and 20% - total concentration of 5e8 platelets/ml) as an initial condition was investigated. The variations delivered AP concentration levels spanned over two orders of magnitude ranging from 5.02e6 platelets/ml to 1e8 platelets/ml, however did not change the scaled AP concentration values.

#### Influence of chemical activation

A closer look at the baseline simulation with 9000 rpm and 2 l/min has shown that the ADP threshold value responsible for the chemical platelet activation was never reached. The highest value within the domain was 1.53e-4 [mol m^-3] which is far below the ADP_t_ value of 2e-3 [mol m^-3]. Running the model without any chemical activation thus produced the exact same results. Reducing the ADPt to 1e-4 [mol m^-3] led to model collapse due to very high source terms of the convection-diffusion equation and a negative volume fraction φ_f_=φ_a_/(φ_a_+φ_n_). To circumvent this problem, we increased the characteristic time of chemical activation t_ADP_ from 1s to 100s. The simulation produced an extreme increase of activated platelets in region 2 (fore bearing) and did not affect the other regions, see Supplementary Figure S6.

## Discussion

The aim of this work was to develop a computationally efficient thrombus model in rotary blood pumps that can be easily implemented and adapted with current CFD solvers.

The model is based on the work of Taylor et al. [11] considering mechanical and chemical activation of platelets and convection-diffusion transport of three species: non-activated platelets, activated platelets and ADP. There are three main differences distinguishing our approach from the model of Taylor et al. First, the species transport equations are uncoupled from the solution of the flow field. The convection-diffusion equations are solved in a domain with the converged velocity field (either steady state or last time step of a transient simulation) as initial conditions. This allows to drastically increase the time-step size from 1e-4 [s] to 1 [s] to solve the convection-diffusion equations resulting in additional computational costs between 10-20 core-hours. Second, the formation of a clot and its influence on the velocity field is excluded for our model and thrombus risk is considered as the concentration of the activated platelet species AP. Lastly, in our model the shear stress tensor including laminar and Reynolds stresses is considered for the mechanical activation (equation 1).

Our model correlated well (R^2^ > 0.94) with the study by Rowlands et al. [16] even considering the fact that aspects as flow rate and rotational frequency were not exactly known in the clinical study or were probably different for each patient.

Analyses of different speeds between 7000 and 11000 rpm and of different flows between 1 and 4 l/min were carried out. Higher rotational speeds cause higher concentrations of AP. The increase in regions downstream of the impeller was greater than in regions upstream of the impeller. This observation can be explained by the fact that the mechanical activation is increasing in the impeller region due to the higher shear stresses at higher speeds and the activated platelets stay in recirculation regions around the diffusor. In low flow simulations, considerably higher concentrations of activated platelets were predicted than in high flow simulations. This is also consistent with the observations of Wu et al. [15]. Low flow operating conditions have also shown to lead to increased hemolysis both in silico [29] and in vitro [30] due to higher recirculation and longer exposure times of red blood cells. Probably a similar effect takes place for the platelets.

Influencing factors of the different components of the accelerated thrombosis model were investigated. The sensitivity analysis of the parameters included in the shear stress power law model showed that only the parameter *α* had a large influence on the activated platelet concentration. This influence, however, applied equally to the entire domain, thus raising the overall activated platelets concentration level to a similar extent. Hence, the relative contribution of different pump compartments towards thrombus formation stayed the same. In addition, the influence of turbulence effects on the scalar shear stress was investigated. Compared to the laminar approach, the activated platelet concentration increased especially downstream of the impeller blades, a region where one would expect a high contribution of turbulent stresses. It has to be noted, that a Reynolds-averaged Navier–Stokes equations (RANS) turbulence model has been used. Scale resolving models such as Large Eddy or Detached Eddy Simulations might produce different contributions of turbulent stresses, potentially higher than the ones of the RANS model. Changes of the initial ratio of activated to non-activated platelets raised the absolute value of AP but did not affect the relative contribution of different pump components. Lastly, it was shown that chemical activation did not have any influence on the results as the activation threshold value ADP_t_ was never reached. Lowering this threshold value (and increasing the t_ADP_ from 1s to 100s to achieve stable simulations) led to a drastic increase of activated platelets in the fore bearing region (Volume 2) but did not affect other parts of the pump. Such a “switch-like” behavior is one of the main weaknesses of the model and a continuous relationship between ADP concentration and activated platelets, e.g. through a Hill-like function, should be implemented in the future.

Our approach compares to the existing thrombogenicity models in the following ways. As outlined in the introduction, PAS, which is easy to implement, is often taken as a global measure to compare different designs with each other and does not focus on different regions of the devices [31]. A recent implementation by Chiu et al. [10] used a particle approach and computed a probability density function from the stress accumulation of particles, compared two VAD designs with each other and also focused on the probability density functions in the different regions of the pumps. The global in silico comparison (device 1 vs. device 2) was confirmed with in vitro and in vivo experiments, however it remains unclear to what degree the localized thrombus risk can be trusted upon. The particle based approach has, in general, limitations regarding correct stress accumulation in narrow gaps of the pump, an observation that has also been made for hemolysis models [9].

The model by Wu et al. [14] provides more information on thrombus mechanisms by consideration of 10 instead of just 3 biochemical species. It clearly gives a more thorough description of thrombosis than our model at a cost of higher parameter uncertainty (3 parameters such as the deposition rate of platelets for the materials that come into contact with the blood must be determined experimentally) and higher computational costs of 200 core-hours vs. 20 core-hours.

A third modelling approach is a recent study by Dai et al. [32] employed a two-phase approach of two fluids with identical rheological properties and defined thrombosis risk regions as regions of low washout. Although computationally cheap, the approach is purely phenomenological and depends on definitions of “washout” regions. Even if such thresholds are identified for one pump geometry (with existing in vitro data), there is no guarantee that these values will hold for other device designs. Lastly, the model of Taylor et al. [11] was developed and experimentally validated in a backward facing step and, as already explained in the introduction, would exhibit extremely high computational costs when implemented in a rotary blood pump environment.

All things considered, we believe that our model provides a promising trade-off between complexity and computational costs. It includes a higher degree of mechanistic detail compared to the PAS approach but does not include as many mechanisms as the model by Wu et al. [24]. The approach of uncoupling the flow and the species transport together with neglecting a physical presence of a thrombus drastically reduces computational costs but still allows a reliable comparison with clinical data. Using a steady state (or transient averaged) solution of the velocity field neglecting dynamic flow features potentially works for continuous flow pumps. New generations of VADs include additional pulsatility, such as the Lavare cycle of the HVAD and the artificial pulse of the HeartMate III [33]. To consider the influence of pulsatility one could neglect the uncoupling and combine flow field calculations with the species transport. As an alternative, considering two species transport simulations, one with a transient averaged velocity and one with the root mean square velocity might be a possible approach. As already mentioned above, a more detailed representation of chemical activation (Hill-like activation and consideration of further chemical species) might be a next possible steps. In addition, influence of the coagulation cascade and the influence of artificial materials on platelet activation could be included, like in the model by Mendez et al. [34].

As it is extremely difficult to obtain experimental data on thrombus formation in medical devices, all computational studies should be handled with care. To the best of our knowledge, the study by Rowlands et al. [16] is the only study that provides quantitatively reliable data on thrombus deposition, although in a pump that is continuously substituted by new generation VADs. Still, we cannot know for sure if the model performs equally well in other pump geometries. Our exploration of the Medos DP3 centrifugal blood pump, see Supplementary Figure S8, showed increased concentration of activated platelets in the bearing region and in the secondary gap which seems to correspond with clinical observations of pump head thrombosis of the DP3 [35]. Nevertheless, it is not appropriate to consider this observation a validation of the computational model.

## Conclusion

Within this study we presented a computationally efficient model of thrombosis for CFD simulations in rotary blood pumps. The model exhibits equal or better accuracy and provides lower computational costs than common state of the art approaches. The implementation of the model in the FDA Round Robin geometry within the commercial solver ANSYS CFX has been included under (https://doi.org/10.5281/zenodo.5116063) for further testing and expansion.

## Acknowledgments

The authors thank Dr. Bente Thamsen for the reconstructed HeartMate II geometry.

