## Supplementary Information for "An accelerated thrombosis model for computational fluid dynamics simulations in rotary blood pumps"

***Correspondence:**

### Supplementary methods

#### *Model structure*

To get an overview of how all parameters and equations of the model are related to each other, the following Figure S1 is helpful. It shows the structure of the accelerated thrombosis model as well as the initial model developed by Taylor et al.:

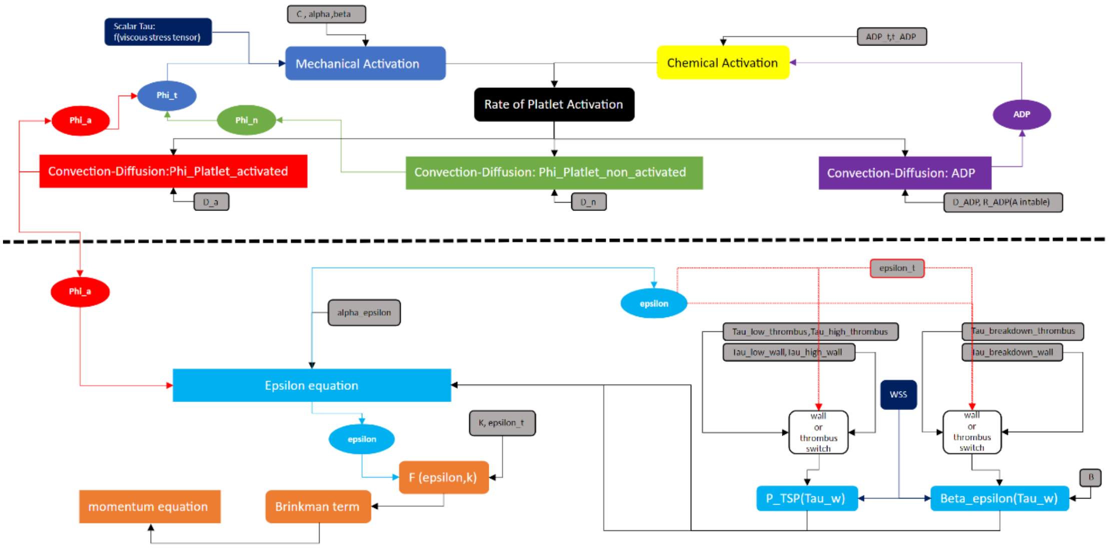


*Figure S1: Overview of all equations and parameters involved in the Taylor model. Above dashed line: Structure and parameters of the accelerated thrombosis model*

Equations are marked with square fields, species with oval fields and influence factors and parameters with rounded rectangular fields.  Above the dashed dividing line in the middle of are the three convections diffusion equations and their influencing parameters. This area is therefore also the area of the accelerated thrombosis model.

#### *Bearing representation*

In the Figure S2 the bearing is shown oversized to visualize more clearly the way it is constructed. In the real geometry the distance between the two bearing shells is 0.05 mm and the bearing shell diameter is approx. 3 mm.  The rear bearing is constructed in the same way.

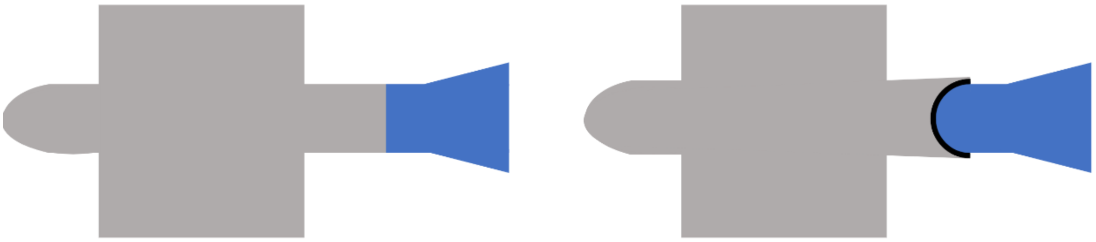

**a)** **b)**

*Figure S2: Cross-section of front area of Heartmate 2 (a) and of Heartmate 2 with modelled front bearing (b)*

### Supplementary results

*Operating point analysis*

The correlation between numerical and clinical data for different rational speeds and different flow rates is shown in Figure S3.

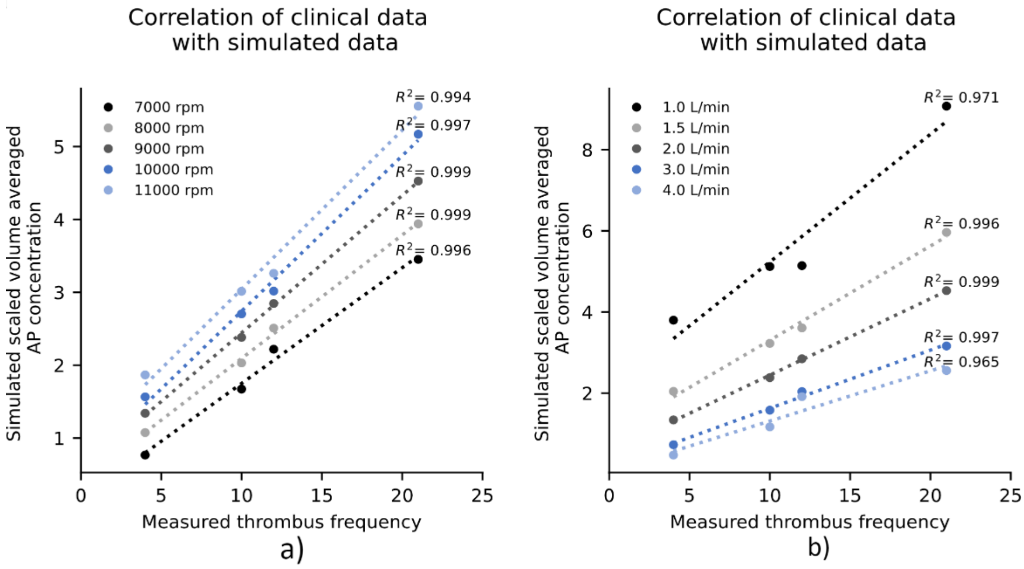


*Figure S3: Correlation between normalized AP platelet concentration and measured thrombus frequency from the Rowlands et al. study for left – rpms ranging from 7000-11000 and right – flow rates ranging from 1-4 l/min*

*Shear stress power law model*

Figure S4 again looks at the distribution of the AP concentration on the surface. The left side of the figure shows the results of the $\alpha$ perturbation simulations with the scale that has been used for the AP concentration contour plots in the main text. An adjustment of the scale for each of the three cases is done on the right side of the figure. The maximum value of the scale is adjusted in such a way that a similar distribution is obtained as in the baseline case with baseline $\alpha$ value. Changing the α value results in different AP concentration levels, but does not change the final output of the simulation, because by adjusting the scale, very similar contour plots can be created on the surface.

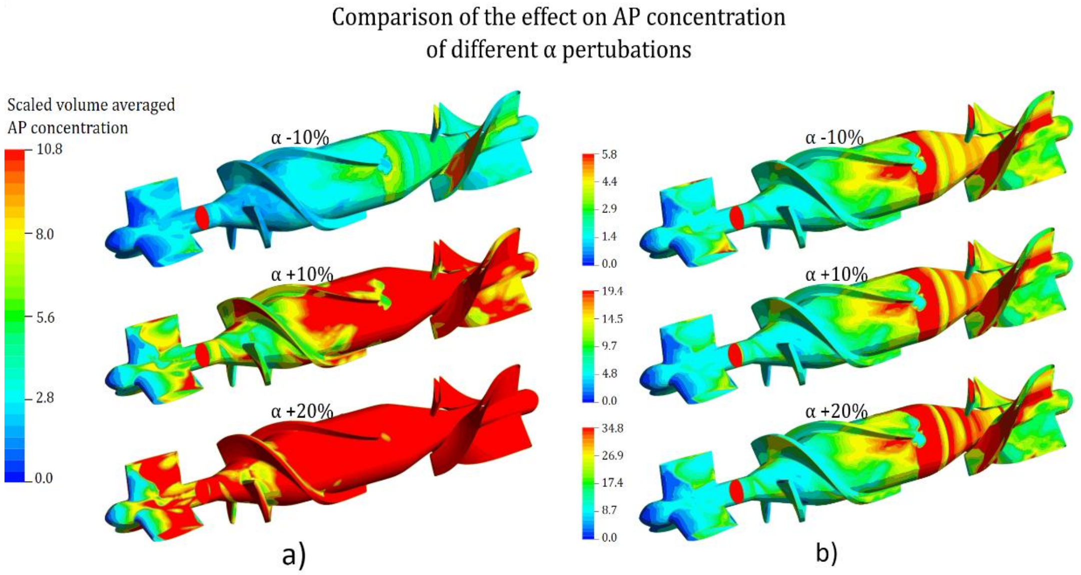


*Figure S4: Comparison of different simulations with varying α parameter using contour plots of the AP variable with same scale (left) and different scales(right).*

*Influence of turbulence modeling*

The effects of the turbulent approach on contour plots and volume renderings are shown in Figure S5. On the left side of the figure, the images of the baseline simulation with the laminar approach are shown for comparison purposes. The scale of the contour plot is again the same as described before and applies for both the laminar and turbulent contour plot. On the right-hand side of the figure the results for the turbulent case are shown. The volume rendering is created with the same threshold value as in the laminar case. This shows that after the bladed passage a higher AP concentration level is present in the entire flow due to platelet activation through the turbulent Reynolds stresses.

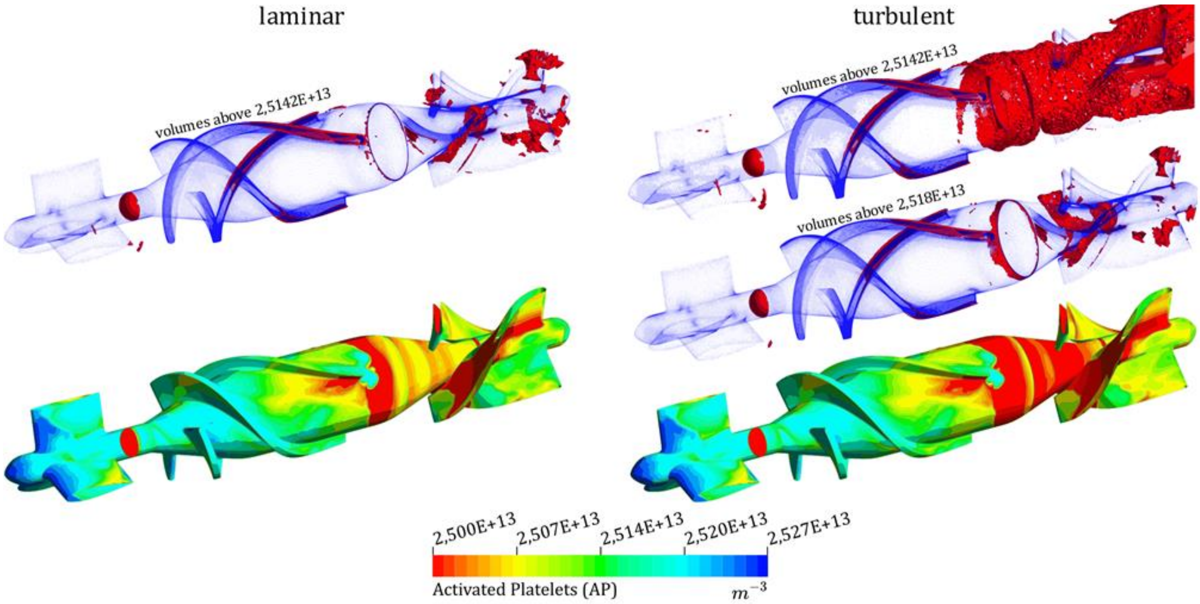


*Figure S5: Comparison of laminar and turbulent shear stress simulations using contour plots and volume renderings of the variable Activated Platelets.*

*Influence of chemical activation*

The results of the simulation with effective chemical activation ($ADP_t$ = 1e-4 [mol m^-3] and activation $t_{ADP}$ =100s) can be seen in Figure S6. The bar chart on the left shows that the simulation with the modified chemical activation produces substantially higher AP concentration values in region 2. On the right side of Figure S6, it can be seen from the contour plots that the AP concentration values in the regions of the bearings of the modified simulation are substantially higher than in the baseline simulation. Furthermore, the values in the areas around the bearings are also slightly elevated. The remaining regions remain unaffected by the change.

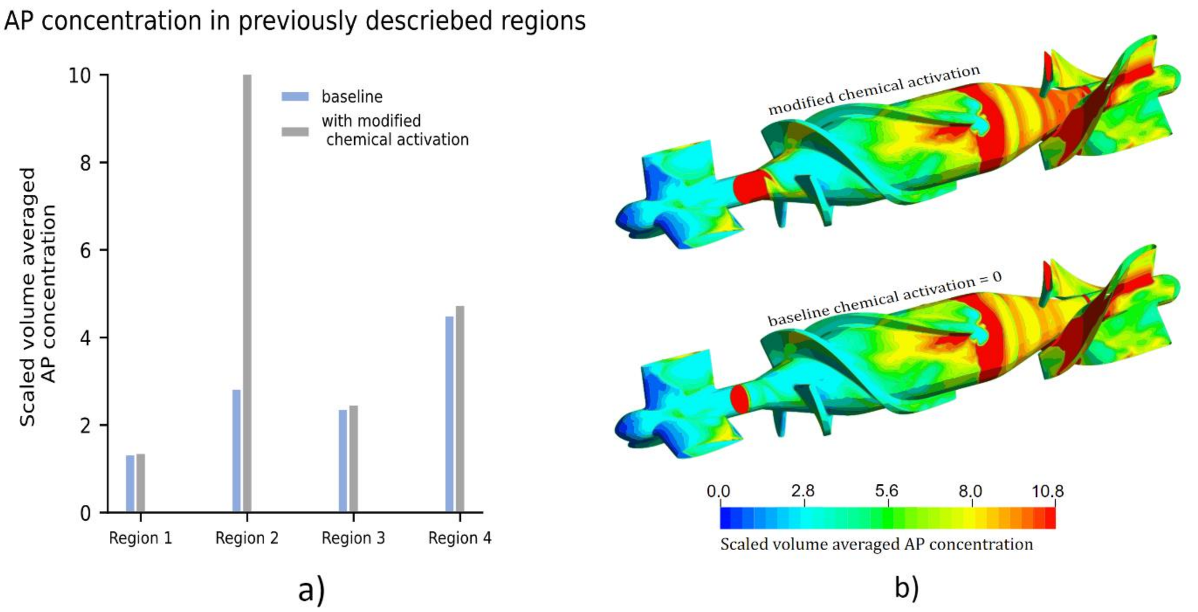


*Figure S6: Comparison between the baseline simulation, which showed no chemical activation, and a modified variant of chemical activation using the variable AP in a bar chart and a contour plot*

*Accelerated thrombosis model with DP3*

In addition, the Medos DP3 geometry from the study by Groß-Hardt et al. [1] was taken to see how the model performs in an extracorporeal centrifugal pump. The meshed geometry is shown in Figure S7. The similar meshing strategy as was used in the HM2 geometry resulted in a mesh of approximately 7 million elements. To see how the model performs in another pump geometry, the accelerated thrombosis model was applied to the DP3 geometry at 5000 rpm and 1.5 l/min.

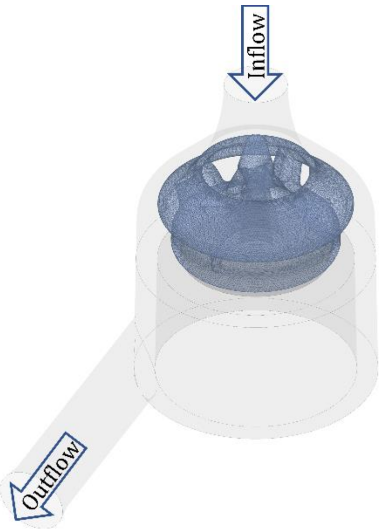


*Figure S7: Meshed geometry of the Medos DP3 pump*

Figure S8 – left, illustrates the AP concentration distribution over the simulation domains on the left side. The AP concentration is initially low in the inlet section and then increases abruptly from the rotor domain onwards. The concentration is highest in the Secondary Gap region. In Figure S8 – right, it is possible to identify that the volumes with the highest AP concentration are located exactly at the bearing of the impeller.

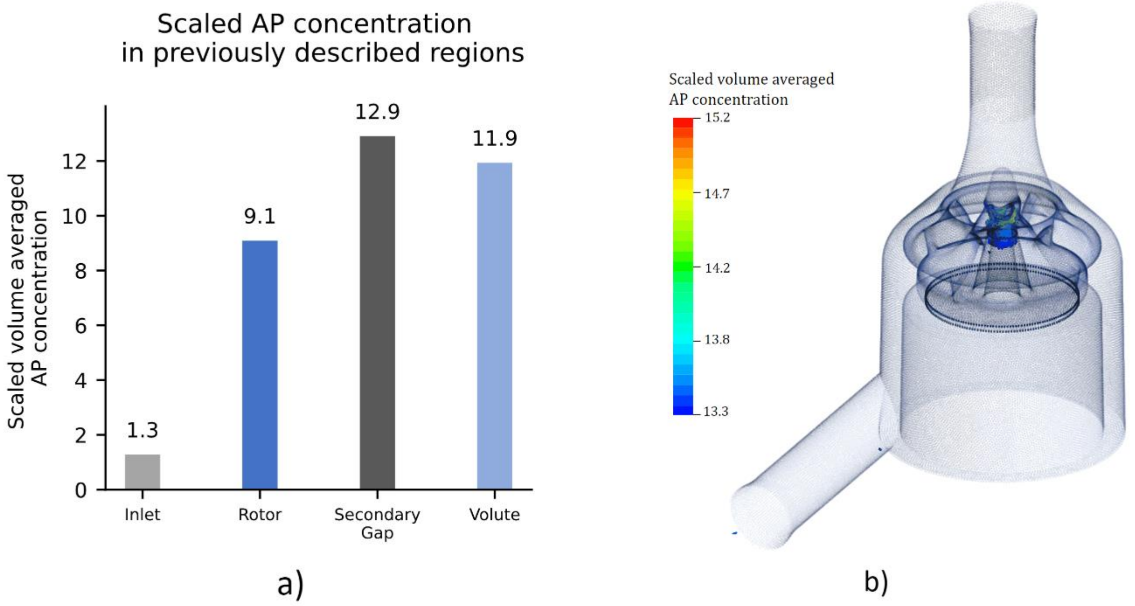


*Figure S8: Application of the accelerated thrombosis model to DP3 geometry at 5000rpm and 1.5 L/min and the resulting AP concentration*
